# Conformational Dynamics of the Nuclear Pore Complex: Insights from Elastic Network Models and Coarse-Grained Simulations

**DOI:** 10.1101/2025.05.28.656622

**Authors:** Hengyue Wang, Guanglin Chen, Zhiyong Zhang

## Abstract

The nuclear pore complex (NPC) serves as the primary channel regulating nucleocytoplasmic transport in eukaryotic cells. In recent years, atomic-resolution structural models of both the constricted and dilated states of the NPC central core have been resolved; however, the molecular mechanisms underlying the transition between these two states remain unclear. This study employed elastic network models (ENM) to identify the first three low-frequency collective motion modes of the NPC scaffold structure, revealing the intrinsic relationship between these modes and the two-state transition of the NPC scaffold. Molecular dynamics (MD) simulations were further utilized to achieve dynamic conformational transitions between the constricted and dilated states of the NPC scaffold, providing detailed insights into the dynamics of this process. Regarding the FG repeat sequences within the nuclear pore channel, simulations based on the inner ring heterotrimeric FG-Nups revealed significant differences in their distribution between the constricted and dilated states. Finally, by integrating multi-scale research findings, we propose a potential nucleocytoplasmic transport regulatory pathway that encompasses both NPC conformational changes and FG barrier dynamic behavior, providing novel support and validation for the dilation model.

## INTRODUCTION

Eukaryotic cells store their genetic material within the nucleus and segregate it from other cellular components through a double nuclear membrane. This compartmentalization enables cells to perform complex and specialized biological activities while simultaneously presenting challenges for material exchange between the nucleus and cytoplasm. The vast majority of material exchange occurs through channel structures in the nuclear membrane—the nuclear pore complexes (NPCs). The transport processes governing molecular entry and exit from the nucleus determine the subcellular localization of numerous biological macromolecules, including transcription factors, and serve as the foundation for gene expression regulation, cell division, and other critical cellular functions[1].

The nuclear envelope consists of inner and outer lipid bilayers, with NPCs embedded at nuclear pores where the inner and outer membranes curve and fuse. Human NPCs can be conceptualized as hollow cylindrical structures with an outer diameter of approximately 1200 Å, a height of approximately 800 Å, and a total mass of approximately 120 MDa[2, 3]. NPCs can be divided into three components: the central core (comprising the inner ring, cytoplasmic ring, nuclear ring, and luminal ring) that associates with the nuclear membrane and forms a diffusion barrier; eight filaments (cytoplasmic fibers) that connect to the core and extend into the cytoplasm; and a basket-like structure (nuclear basket) formed by eight filaments on the nuclear side. All components exhibit eight-fold rotational symmetry around the central channel axis[2].

Structural biology studies, employing techniques such as cryo-electron microscopy, have resolved atomic-level structures of the central cores of NPCs from multiple species, particularly human NPCs[4–7]. These high-resolution structures have revealed the precise spatial configuration of NPCs as hollow cylinders and clarified the spatial distribution relationships among constituent substructural units. NPCs comprise approximately 1000 nucleoporins (Nups), encompassing about 34 distinct Nup types encoded by the human genome. Overall, Nups can be classified into three categories: ① transmembrane Nups, responsible for anchoring NPCs to the nuclear membrane and maintaining structural integrity[8, 9]; ② structural Nups (or scaffold Nups), serving as the primary framework of NPCs and conferring fundamental mechanical stability and spatial structure[10]; and ③ FG-Nups containing phenylalanine-glycine repeat sequences, which form highly dynamic selective barriers within the central channel and constitute the core determinant of nucleocytoplasmic transport selectivity[11–15]. Through ordered assembly, these three categories of Nups collectively construct the overall NPC structure and functionality characterized by typical eight-fold symmetry[8, 16–20].

In situ studies have demonstrated that the diameter of the NPC central channel and its spatial configuration undergo significant changes in response to different physiological conditions or environmental variations[14, 21–26], suggesting that NPCs possess high conformational plasticity in vivo. The static conformations of human NPCs revealed by atomic-resolution structures fail to provide insights into these dynamic processes[27, 28]. Computer simulations, particularly molecular dynamics (MD) methods, have emerged as powerful tools for exploring the dynamic behavior of FG-Nups and molecular transport mechanisms[6, 7, 29–34]. Existing simulations have primarily focused on the interactions between FG-Nups and transport receptors[35, 36], conformational fluctuations of FG repeat sequences[31, 37, 38], and how these factors influence selective barrier mechanisms[7, 32, 39–43]. However, most related simulation studies have employed fixed NPC scaffold structures, with limited attention to the dynamic changes in scaffold architecture and their effects on FG-Nup distribution and functional states.

Based on the current research landscape, this study aims to integrate elastic network model (ENM) analysis with molecular dynamics simulations, focusing on the dynamic changes in NPC scaffold structure and their regulatory effects on FG-Nup distribution characteristics and nuclear pore selective barrier formation. Specifically, we will systematically analyze the low-frequency collective motion modes of the NPC scaffold, investigate their relationship with nuclear membrane morphological changes and nuclear pore channel diameter variations, and further elucidate the molecular mechanisms by which scaffold structural dynamics regulate the spatial distribution of FG repeat sequences. This approach will provide novel structural biology and computational simulation evidence for understanding the dynamic nature of NPC selective barriers.

## MATERIALS AND METHODS

### The human nuclear pore complex

The human nuclear pore complex scaffold structure (PDB ID: 7r5k[7] and 7r5j[7]) comprises four components: the inner ring, luminal ring, cytoplasmic ring, and nuclear ring. Here, 7r5k represents the constricted state NPC, while 7r5j represents the dilated state NPC. In this study, to simplify the simulation system, the luminal ring component was uniformly removed.

### Construction of the coarse-grained simulation system

We constructed the coarse-grained NPC scaffold structure and surrounding nuclear membrane pore system following essentially the same procedures as previous studies[7]. However, when completing the missing residues between Nup protein chains in the NPC scaffold structure model, this study employed the IDRWalker[44] tool. The equilibration simulation parameters were also largely consistent with previous studies, conducting 10 ns equilibration simulations with a 2 fs time step.

### Elastic network model

This study employed the basic anisotropic elastic network model (ANM)[45] algorithm. Unlike directly using PDB database structures for ENM calculations, to eliminate potential non-physical conformations in the initial structures, this study performed ENM calculations on the constricted and dilated conformations obtained after short-term equilibration MD simulations.

The dimension of the Hessian matrix is 3N_r_x 3N_r_, with its scale growing quadratically with the number of nodes N_r_ . In this study, the NPC scaffold system contains N_r_ = 516,192 amino acid residues, corresponding to a Hessian matrix with (3N_r_)^2^ = 2,398,087,627,776 elements (approximately 2.4 trillion), far exceeding the computational capacity of eigenvalue solver libraries such as LAPACK and ARPACK.

Based on the eight-fold rotational symmetry of the NPC scaffold structure, this study introduced periodic boundary condition constraints into the classical ANM algorithm, decomposing the complete system into 8 equivalent symmetric units (subunits) for independent calculation, ultimately reconstructing the overall motion modes. This strategy reduced the number of nodes to N_r_⁄8 = 64,524, with the corresponding Hessian matrix elements reduced to (3N)^2^ = 37,470,119,184(approximately 37.4 billion), significantly lowering computational complexity.

The NPC eight-fold rotational symmetric structure is parameterized as 8 isomorphic units (numbered 1-8) distributed in a ring configuration, where each unit i establishes spatial proximity relationships with adjacent unit j according to the principle of cyclic symmetry, with j=(i mod 8)+1 (i=1,…,8). The system constructs eight groups of adjacent unit pairs (1,2),(2,3),…,(7,8),(8,1), and within each unit pair (i,j), amino acid residues participate in local minimum distance calculations and corresponding vector solutions. For each unit pair, an inter-residue distance dataset DISTANCE = |A1B1|, |A1B2|, |A2B1| and its corresponding vector set VECTOR = A1B1, A1B2, A2B1 are constructed, where Am and Bn represent two equivalent conformational states of residues within symmetric units (m,n in 1,2). The minimum distance rABmin and its corresponding vector AB are selected from these datasets as input parameters for ANM calculations, and finally, the eigenvectors from the 8 symmetric units are integrated to reconstruct the collective motion modes of the complete NPC scaffold structure.

### Two-state transition simulations with PLUMED

Due to the large size of the simulation system, the timescale of conventional molecular dynamics simulations cannot capture the transition process between the two NPC states. Therefore, this study incorporated biasing potentials during simulations, employing the PLUMED[46] plugin to perform nuclear pore complex two-state transition simulations, and analyzed the simulation results using alignment error analysis.

First, in the PLUMED plugin, all protein backbone particles (BB) within each subunit of the NPC scaffold were defined as a GROUP. The center of mass of each GROUP was then calculated, and DISTANCE was defined as the distance between the centers of mass of two centrally symmetric GROUPs. From the NPC scaffold structural models in constricted and dilated states, the standard DISTANCEs for the constricted and dilated states were measured. Finally, the MOVINGRESTRAINT function in the PLUMED plugin was invoked to perform NPC two-state transition simulations in the NPT ensemble: using the C-rescale pressure regulator and V-rescale thermostat, with the pressure coupling time constant set to 40 ps. Within 100 ns, using a 20 fs time step and a force constant of KAPPA = 10,000 kJ·mol_⁻_¹·nm_⁻_², the current DISTANCE was stretched or compressed to the corresponding standard DISTANCE to achieve NPC scaffold two-state transition simulations. This study conducted a total of six simulation sets, with three sets in each direction.

### Aligned error

This study treats each Nup as the basic unit for AE analysis. First, the RMSD between the initial and final states of Nup-A is calculated. Since RMSD calculation requires finding the alignment operation that minimizes RMSD, this study extracts this alignment operation separately and applies it to Nup-B, calculating the RMSD between the initial and final states of Nup-B under this operation. This RMSD is defined as the AE from Nup-A to Nup-B. Similar to how RMSD magnitude can measure the degree of internal structural change within a single Nup, this study expects to reflect the magnitude of relative positional changes between two Nups through AE values.

Since no long-range interactions exist between proteins, this study only considers AE between physically proximate Nups and sets a cutoff distance of 10 nm. Finally, only AE between two Nups whose center-of-mass distance is less than the cutoff distance in the NPC constricted state is calculated. Other AE values excluded by the cutoff distance are set to -5 for visualization purposes.

The NPC scaffold structure possesses eight-fold rotational symmetry. For AE calculations between any two Nups, all symmetrically equivalent positions must be traversed and arithmetically averaged. To simplify this process, this study employs an algorithm similar to that used in ENM analysis: only considering AE between Nups in adjacent subunits, then arithmetically averaging the minimum AE values from the 8 groups of adjacent units. Finally, this study obtained the average AE matrix between all Nups in the NPC two-state transition simulations.

### Simulations of the inner-ring FG-Nup system

The inner-ring FG-Nups primarily comprise three types of Nups: Nup54, Nup58, and Nup62. These Nups exist in 4 copies within each subunit’s inner ring, resulting in a total of 32 copies (4×8) in the complete NPC scaffold inner ring. This study extracted all copies of these Nups from the NPC scaffold inner ring as the non-FG repeat sequence portion of the inner-ring FG-Nups. Based on the positions of FG repeat sequences within the FG-Nups, this study used IDRWalker[44] to randomly generate the spatial distribution of FG repeat sequences.

To better observe the dynamic conformations of FG repeat sequences, this study employed the Martini3 coarse-grained force field[47] with modified parameters[48]. The Martinize.py script was executed to individually coarse-grain each inner-ring FG-Nup protein chain using the Martini3 coarse-grained force field. Parameter settings were as follows: all chain termini were set to uncharged states; protonation states adopted default settings. Since the research subjects are intrinsically disordered proteins (IDPs) rather than conventional ordered proteins, this study did not use the DSSP[49] algorithm to assign secondary structures but uniformly set the secondary structure to "C" (coil), representing disordered regions; elastic networks were also not employed to maintain tertiary structure.

Although the modified Martini3 parameters can effectively simulate the dynamic behavior of IDPs, they correspondingly cannot properly handle protein interactions and dynamic behavior in conventional ordered regions[50]. Therefore, throughout the entire simulation process (including energy minimization), this study applied position restraints to the protein backbone beads (BB) of the non-FG repeat sequence portions of FG-Nups, with a force constant of 1000 kJ·mol_⁻_ ¹·nm_⁻_ ².

This study employed position restraint settings in GROMACS[51]: cylindrical restraints (Cylinder) constraining the XY directions and layer restraints (Layer) constraining the Z direction, with the centers of both the cylinder and layer set at the center of mass of the non-FG repeat sequence portion of the inner-ring FG-Nups. Both the cylindrical radius and layer height were set to 0.01 nm to implement restraining forces directing all FG repeat sequences toward the nuclear pore center. The restraining forces were configured in 5 levels: (0.1, 0.05, 0.01, 0.005, 0.001) kJ·mol_⁻_ ¹·nm_⁻_².

Energy minimization was performed using steepest descent and conjugate gradient methods to eliminate steric clashes in the initial model. Subsequently, equilibration was conducted in the NVT ensemble: temperature was maintained at 310 K through the Berendsen thermostat. The Verlet neighbor search algorithm was used to update particle neighbor lists, with both cutoff distance and update frequency automatically optimized by GROMACS[51]. The cutoff radius for Lennard-Jones and Coulomb forces was set to 1.1 nm, with potential energy zero-point correction achieved through the Verlet displacement potential corrector. Short-term equilibration simulations were run for 10 ns with a 2 fs time step, without position restraints on FG repeat sequences.

The thermostat was changed to V-rescale for FG repeat sequence position-restrained simulations in the NVT ensemble, conducted in three steps. In the first step, the force constant for FG repeat sequence position restraints was set to 0.1 kJ·mol_⁻_¹·nm_⁻_², and simulations were run for 100 ns with a 20 fs time step to aggregate FG repeat sequences at the nuclear pore center. In the second step, based on conformations where the majority of FG repeat sequences had aggregated at the center, the force constant was modified to (0.05, 0.01, 0.005, 0.001) kJ·mol_⁻_ ¹·nm_⁻_², and 4 groups of simulations were conducted for 100 ns each with a 20 fs time step to observe the dynamic conformations of FG repeat sequences under different force constant magnitudes. In the third step, after the FG repeat sequence distributions under these 4 force constants reached stable states, position restraints on FG repeat sequences were removed, and 4 groups of simulations were conducted for 100 ns each with a 20 fs time step to observe whether the FG repeat sequence distributions formed through position restraints could exist stably.

## RESULTS AND DISCUSSION

### Periodic boundary optimization verification

To verify the necessity of periodic boundary condition optimization, this study conducted comparative analysis of ENM calculation results for dilated NPC subunits under two conditions: with and without periodic boundary optimization. A and C in Figure 1 show the first eigenvector distribution of isolated subunits, while B and D in Figure 1 present the results with periodic boundary optimization considered as shown in the first eigenmode figure.

**Figure 1.**
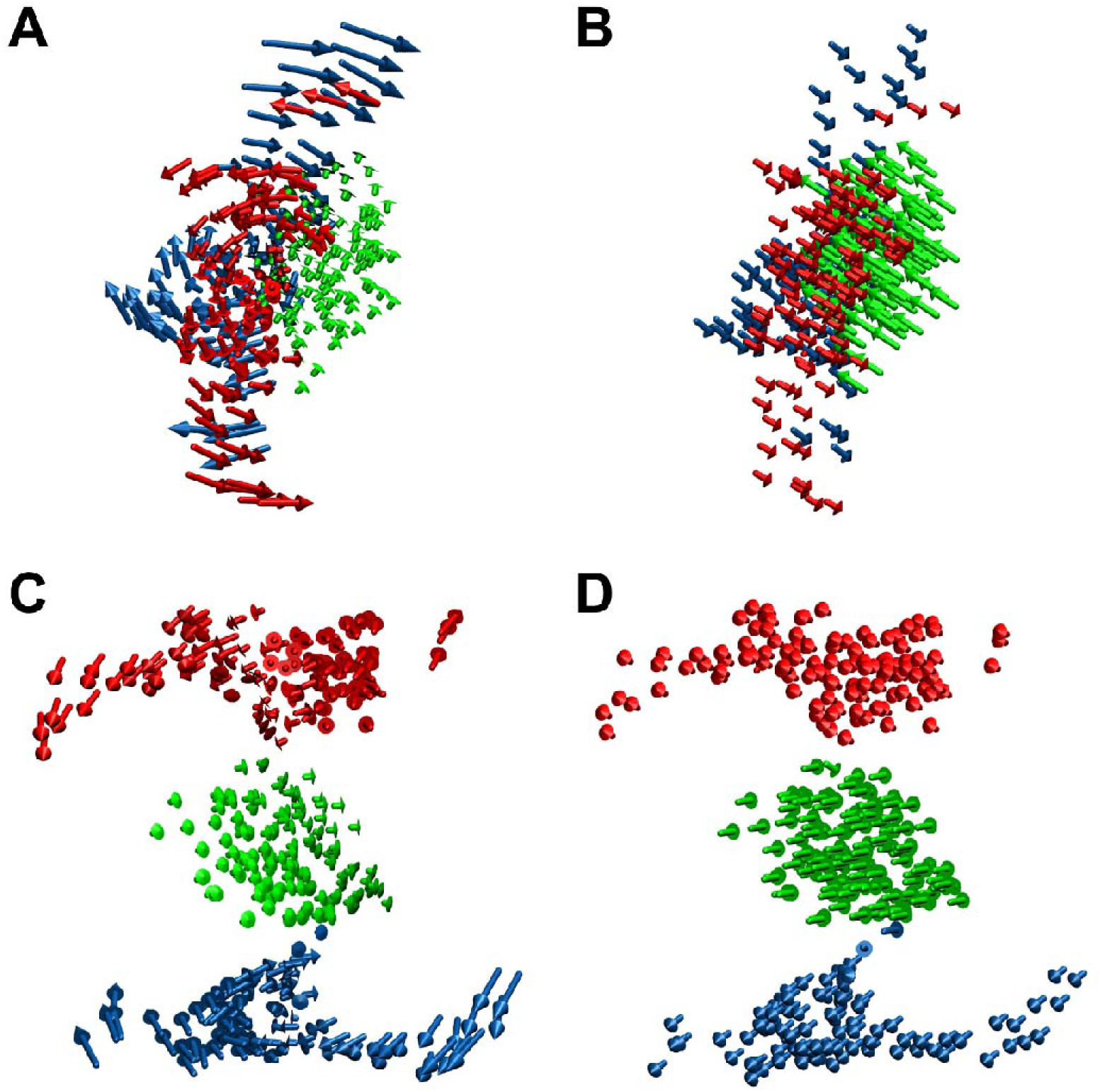
: Comparison of eigenvectors for individual subunits with and without periodic boundary condition optimization. The systems in the figure are all individual subunits of dilated state NPCs. Red represents the cytoplasmic ring, green represents the inner ring, and blue represents the nuclear ring. (A) Without periodic boundary condition optimization, top view. (B) With periodic boundary condition optimization, top view. (C) Without periodic boundary condition optimization, side view. (D) With periodic boundary condition optimization, side view.

Comparative analysis reveals that without considering periodic boundary conditions, the magnitude and direction of eigenvectors within each ring exhibit disordered distributions. Particularly noteworthy is the abnormal increase in eigenvector magnitude at structural boundary contact regions in Figure C, reflecting structural instability of boundary nodes due to missing interactions with adjacent subunits. Although the top-view perspective in Figure A shows a common rotational motion trend around the axis for each ring, its specific dynamical significance is difficult to interpret clearly. Compared with results considering periodic boundary optimization, the first eigenvector of isolated subunits fails to exhibit any significant collective motion patterns, demonstrating the importance and necessity of periodic boundary optimization for accurately capturing low-frequency collective motion modes of NPC scaffold structures.

The periodic boundary condition optimization algorithm has inherent limitations, rooted in the fact that the eight-fold rotational symmetric structure of NPCs does not perfectly conform to strict periodic boundary condition requirements. This is specifically manifested when nodes A and B are located in subunit boundary regions, where a 45° angular deviation exists between the connection vector from B to adjacent subunit A and the connection vector from A to adjacent subunit B. However, due to the positive definiteness requirement of the Hessian matrix necessitating matrix symmetry, only one element hAB = hBA can be retained in the matrix to represent the interaction between A and B within subunits. Therefore, two processing standards are formed in the periodic boundary condition optimization algorithm: during the "symmetric unit topological modeling" process, subunits are named using clockwise and counterclockwise directions respectively, enabling ENM calculations to uniformly adopt connections with clockwise or counterclockwise subunits as the computational basis.

Does this difference in selection standards affect the eigenvalues and eigenvectors obtained from ENM calculations? This study performed ENM calculations on the complete NPC scaffold structure using both clockwise and counterclockwise standards. After visualizing the eigenvectors corresponding to the first three low-frequency modes (essential modes) (Figure 2, Figure 3 and Figure 4), we observed that eigenvector amplitudes are essentially consistent, with only overall angular shifts in eigenvector components in the XY plane. It should be noted that regardless of which standard is used for ENM calculations, the obtained results contain certain errors, but these errors are limited to the XY plane and do not affect qualitative analysis of overall NPC motion patterns.

**Figure 2.**
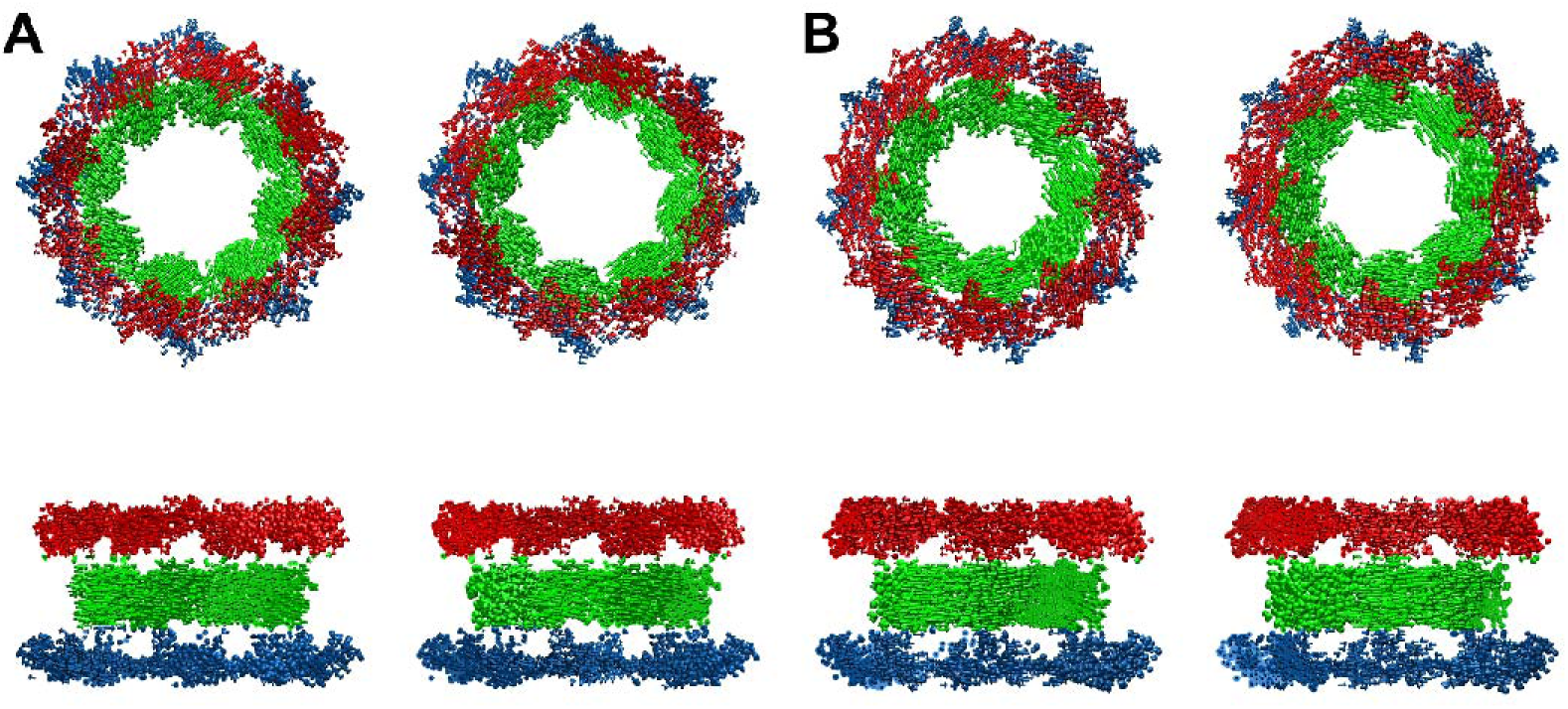
: First eigenvector of the NPC scaffold structure. Red represents the cytoplasmic ring, green represents the inner ring, and blue represents the nuclear ring. The upper panels show top views, and the lower panels show side views. (A) Dilated state, with left and right panels representing the two computational standards in periodic boundary condition optimization, respectively. (B) Constricted state, with left and right panels representing the two computational standards in periodic boundary condition optimization, respectively.

**Figure 3.**
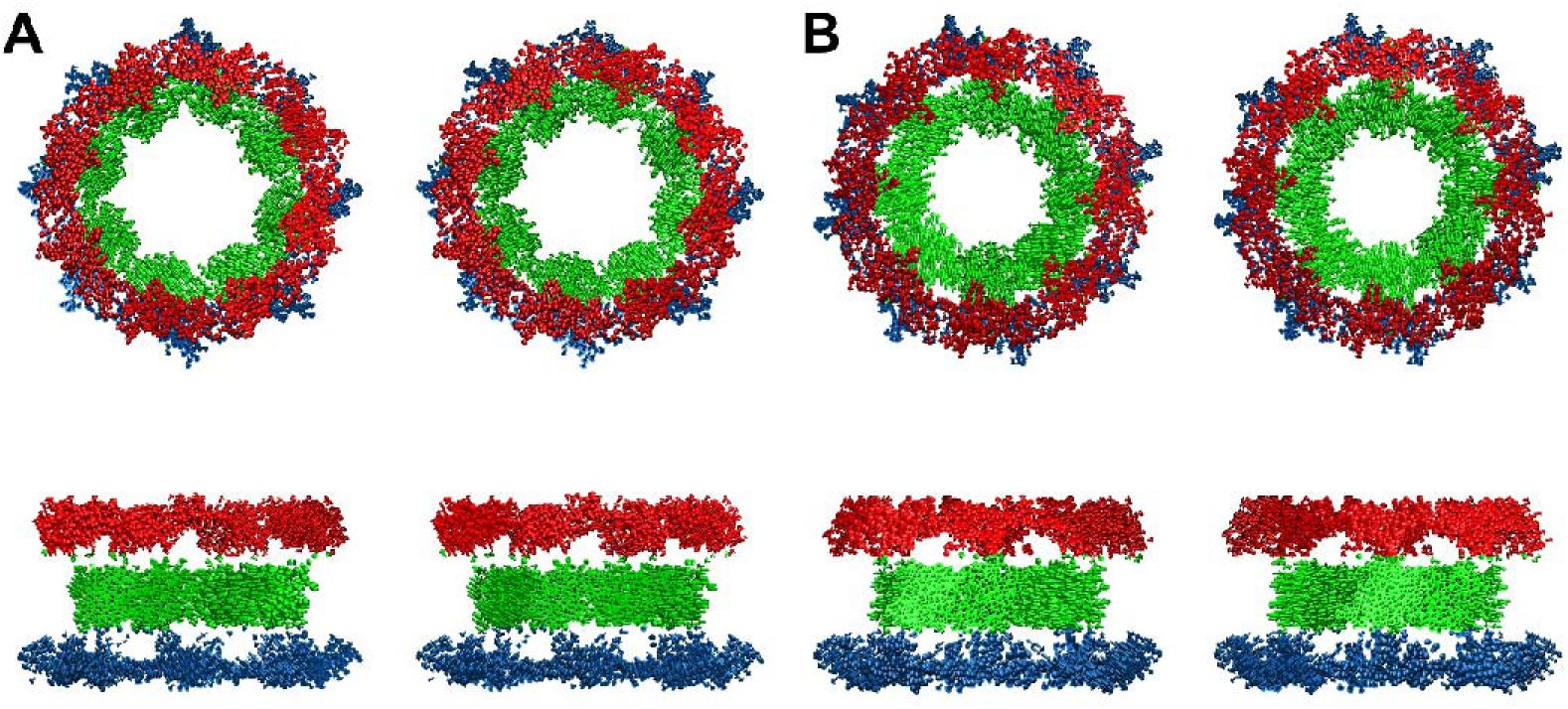
: Second eigenvector of the NPC scaffold structure. Red represents the cytoplasmic ring, green represents the inner ring, and blue represents the nuclear ring. The upper panels show top views, and the lower panels show side views. (A) Dilated state, with left and right panels representing the two computational standards in periodic boundary condition optimization, respectively. (B) Constricted state, with left and right panels representing the two computational standards in periodic boundary condition optimization, respectively.

**Figure 4.**
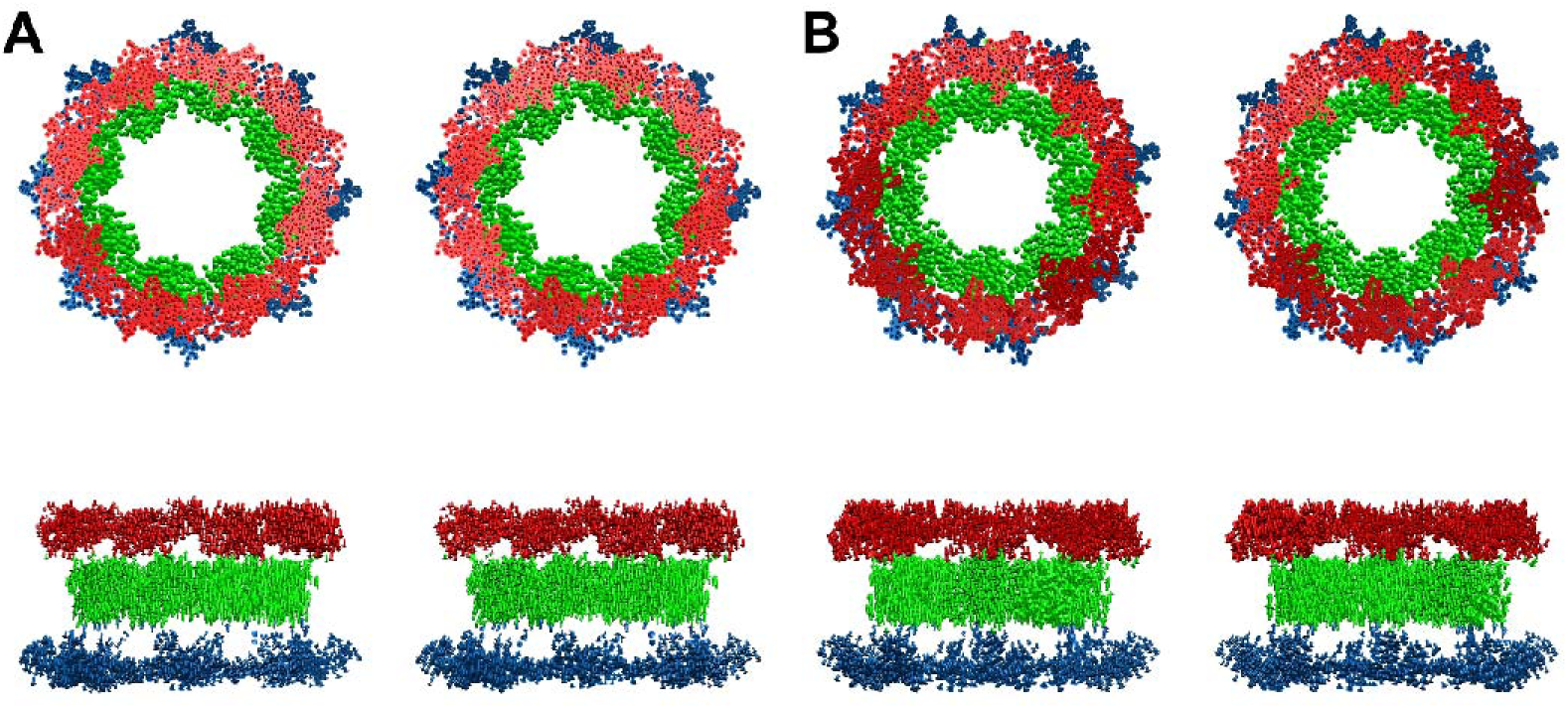
: Third eigenvector of the NPC scaffold structure. Red represents the cytoplasmic ring, green represents the inner ring, and blue represents the nuclear ring. The upper panels show top views, and the lower panels show side views. (A) Dilated state, with left and right panels representing the two computational standards in periodic boundary condition optimization, respectively. (B) Constricted state, with left and right panels representing the two computational standards in periodic boundary condition optimization, respectively.

### Low-frequency collective motion pattern of the first three eigenvectors

Figure 2, Figure 3 and Figure 4 showing the first, second, and third eigenmodes display the spatial distribution of the first three eigenvectors of the complete NPC scaffold structure. Based on the eight-fold rotational symmetry of the NPC scaffold structure, this study defined three orthogonal motion direction basis vectors: (1) axial motion along the Z-axis direction (positive and negative Z directions); (2) radial motion within the XY plane (toward or away from the nuclear pore center); (3) tangential motion within the XY plane (orthogonal to the radial motion direction). Through this decomposition method, the abstract eigenvector directions can be quantified into axial, radial, and tangential components, which together with magnitude parameters describe the collective motion patterns of the system.

Analysis results indicate that the first three eigenvectors exhibit the following common characteristics: using ring structures as boundaries, the collective motion direction and amplitude within each ring of the same subunit are essentially consistent, while significant differences exist between different rings. Among different subunits, the collective motion amplitudes of the same rings are similar, and the motion directions maintain essentially rotational symmetry (symmetry after 45° rotation around the Z-axis) after normalization processing (reversing eigenvectors of some subunits). This consistency in magnitude and symmetry in direction further validates the rationality of periodic boundary condition optimization.

As shown in Figure 2, in addition to the common characteristics of the first three eigenvectors, the first eigenvector exhibits the following properties: within each subunit, the collective motion directions of the two outer rings are essentially consistent and opposite to the inner ring motion direction. Notably, the motion of inner and outer rings is primarily confined to the XY plane, parallel to the plane of their respective rings. As the lowest frequency vibrational mode, this eigenvector reveals the intrinsic mechanical properties of the NPC scaffold structure. First, the opposite motion of inner and outer rings indicates a significant separation tendency. Second, the XY plane-dominated motion pattern suggests that inter-ring interactions are weaker than intra-ring interactions.

These findings are consistent with existing research results. Structural analysis shows that NPC inner and outer rings are connected only through the slender domain of Nup155, with interaction strength far lower than the complex protein networks within rings. Furthermore, the NPC scaffold structure requires nuclear membrane support to maintain stability: its inner surface closely adheres to the nuclear membrane and is anchored through numerous transmembrane Nups. Therefore, although the first eigenvector characterizes the most significant collective motion pattern, this inter-ring separation motion tendency may be suppressed under physiological conditions due to mechanical constraints and regulatory effects of the nuclear membrane.

Unlike the first eigenvector, which is primarily confined to motion patterns in the XY plane, the most significant characteristic of the second eigenvector is that it simultaneously contains Z-axis directional components, manifesting as a coupled mode of axial and planar motion.

First, analyzing the motion components within the XY plane: as shown in Figure 3, the collective motion directions of the inner ring and the two outer rings remain opposite.

Second, in the Z-axis direction, both outer rings exhibit clear axial displacement tendencies, moving away from the inner ring. In contrast, the inner ring shows weaker axial motion tendencies, with the axial motion directions of adjacent subunit inner rings displaying random distribution (either same or opposite). This randomness indicates structural instability of the inner ring, namely weak interactions between inner ring subunits. This study can reasonably infer that inter-outer ring interactions are stronger than inter-inner ring interactions. Finally, analyzing the coupling relationship between axial and planar motion, this coupling is specifically manifested as: contraction of the inner ring within the XY plane couples with outer rings moving away from the inner ring along the Z-axis. Conversely, expansion of the inner ring within the XY plane couples with outer rings moving away from the inner ring along the Z-axis.

Existing research data indicate that outer rings actually exhibit extremely limited structural changes in the XY plane. This is primarily attributed to the dense interaction networks and rigid connection structures between outer ring subunits, whose binding strength is significantly higher than that of the inner ring. Comparison of crystal structures of the two conformational states of NPC scaffold structures in the PDB database shows that outer rings maintain almost completely consistent structural features between constricted and dilated states. In contrast, due to the structural flexibility of Nup155 connecting inner and outer rings, axial displacement of outer rings relative to the inner ring is physically feasible. Therefore, the coupling mode of inner ring planar motion and outer ring axial motion may correspond to real conformational changes: during NPC constriction, outer rings tend to move away from the inner ring; during dilation, they approach the inner ring. However, this inference still requires experimental or MD validation, and no related research has been reported to date.

The third eigenvector presents motion characteristics distinctly different from the first two eigenvectors, primarily manifested as all nodes having motion dominated by Z-axis directional components. As shown in Figure 4, the motion of the cytoplasmic ring, inner ring, and nuclear ring is almost completely restricted to the axial direction, with both outer rings’ axial motion directions still opposite to the inner ring. Additionally, besides axial motion, the nuclear ring exhibits more pronounced radial components away from the nuclear pore center in the XY plane compared to the cytoplasmic ring and inner ring, reflecting structural asymmetry between the outer rings. This asymmetry is also evident in the first two eigenvectors. From a structural composition perspective, this asymmetry stems from differences in outer ring subunit composition: each cytoplasmic ring subunit contains two Y-complexes, while the nuclear ring contains only one, resulting in relatively weaker structural stability of the nuclear ring. Finally, ENM calculation results obtained using two periodic boundary condition optimization algorithm standards show high consistency here, indicating that ENM calculation deviations exist only within the XY plane.

Although the periodic boundary condition optimization algorithm has certain limitations, specifically manifested as potential overall angular shifts in XY plane eigenvector components, this error does not substantially affect the conclusions of this study. Regarding the first eigenvector, this study focuses on examining the overall trends of collective motion between inner and outer rings. While their motion directions in the XY plane are completely opposite, the research does not involve specific motion directions within this plane. Concerning the second eigenvector, the main focus is on exploring the coupling relationship between the radial contraction-expansion components of the inner ring and the Z-axis components of the outer rings. The axial components of the inner ring in the XY plane show clear contraction tendencies under both calculation standards, therefore directional deviations of eigenvectors within the XY plane do not affect the research conclusions. As for the third eigenvector, the analysis primarily focuses on the collective motion trends of the three rings dominated by Z-axis motion, a process unrelated to XY plane components.

In summary, this study draws the following qualitative conclusions. First, the NPC scaffold structure exhibits significant structural instability in the absence of nuclear membrane anchoring. Second, the interaction strengths among structural domains show a gradient distribution: intra-cytoplasmic ring interactions > intra-nuclear ring interactions > intra-inner ring interactions > inter-inner-outer ring interactions. Third, during the constriction process of the NPC scaffold structure (inner ring), the two outer rings exhibit displacement away from the inner ring; during dilation, the two outer rings show motion tendencies toward the inner ring. Among these, the first two conclusions have been supported by multiple structural studies and experimental data, while the third point is proposed for the first time in this study and requires further experimental or molecular dynamics simulation validation.

### Simulation process of two-state transition

During the MD simulation process from dilated to constricted state, the top-view perspective clearly demonstrates the gradual reduction in nuclear pore channel size. The nuclear pore channel presents an irregular circular structure with eight sharp angles distributed at 45° intervals. These sharp angles are formed by gaps between adjacent inner ring subunits and gradually diminish as the nuclear pore channel constricts, nearly disappearing in the final conformation, indicating continuous approach of inner ring subunits during the simulation process. In the simulation from constricted to dilated state, all change trends are opposite to the dilated-to-constricted process.

Through the side-view perspective, richer structural details can be observed during the MD simulation process: The spacing characteristics of the double nuclear membrane, namely the vertical distance between the inner nuclear membrane (adjacent to the nucleoplasm) and outer nuclear membrane (adjacent to the cytoplasm).

Curvature changes at the binding sites between the nuclear membrane and NPC scaffold, quantified using bending angle parameters in this study. As shown in figure 5, angle α represents the cytoplasmic ring-side bending angle, and angle β represents the nuclear ring-side bending angle.

**Figure 5.**
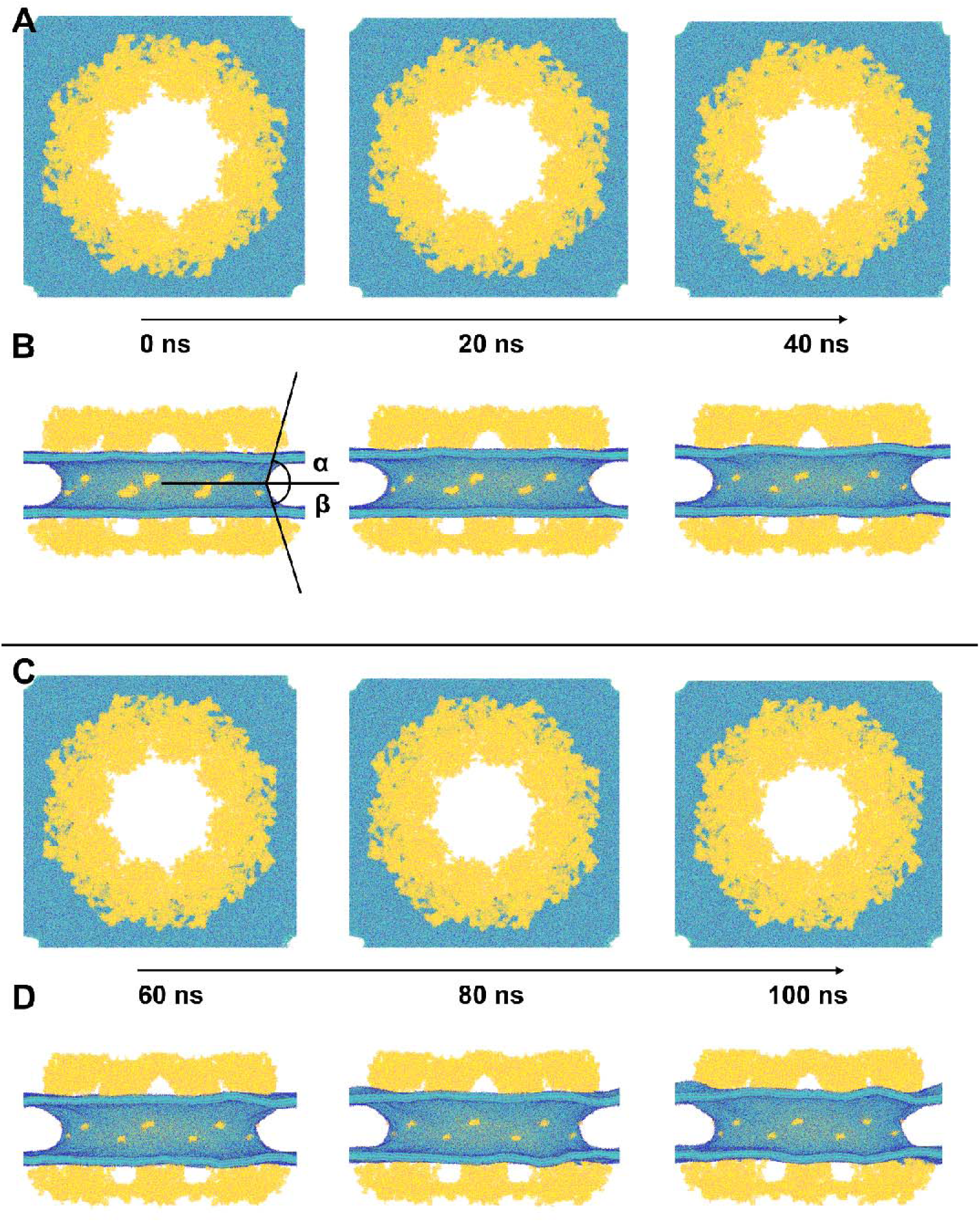
: Process of NPC scaffold structure transition from dilated to constricted state. Six snapshots were selected evenly from the 100 ns simulation process. Yellow represents the NPC scaffold structure, and blue represents the nuclear membrane. (A) System states at 0 ns, 20 ns, and 40 ns, respectively, top view. (B) System states at 0 ns, 20 ns, and 40 ns, respectively, side view. (C) System states at 60 ns, 80 ns, and 100 ns, respectively, top view. (D) System states at 60 ns, 80 ns, and 100 ns, respectively, side view.

The minimum diameter of the nuclear pore channel formed by the nuclear membrane itself. Due to structural symmetry, this minimum diameter typically appears in the nuclear pore center region.

In the side view, the yellow regions within the blue nuclear membrane represent the membrane-anchoring domains of inner ring transmembrane Nups; above the nuclear membrane is the cytoplasmic ring of the NPC scaffold; below the nuclear membrane is the nuclear ring. Although the inner ring structure cannot be directly observed, since it is stably anchored to the nuclear membrane through transmembrane Nups, changes in inner ring dimensions can be indirectly reflected through variations in nuclear membrane channel diameter.

During the simulation process of dilated-to-constricted state transition, the spacing between the double nuclear membranes shows a gradually increasing trend. The nuclear pore diameter formed by the nuclear membrane alone decreases accordingly, consistent with the trend of inner ring subunits approaching each other. Measurement results indicate that bending angles on both cytoplasmic and nuclear ring sides show increasing trends, with the nuclear ring-side bending angle increase significantly greater than that of the cytoplasmic ring side. The increase in double nuclear membrane spacing combined with increased bending angles causes the two outer rings to gradually move away from the inner ring in the Z-axis direction. Additionally, observational data show that neither the diameter nor morphology of the two outer rings exhibit significant contraction or expansion phenomena similar to those observed for the inner ring from the top-view perspective.

During the simulation process of constricted-to-dilated state transition, all change trends are opposite to the dilated-to-constricted process. These observational results mutually validate the conclusions drawn from ENM analysis. During the two-state transition process, the magnitude of changes in the two outer rings is significantly smaller than that of the inner ring, indicating that the interaction strength within outer rings is markedly higher than within the inner ring. Most importantly, the coupling mode of motion between the two outer rings in the Z-axis direction and the inner ring in the XY plane completely matches the predicted results from ENM analysis: outer rings move away when the inner ring contracts, and approach when the inner ring expands. This finding strongly validates the reliability of ENM analysis conclusions.

### Aligned error in the two-state transition

Due to the original numbering rules in the PDB database that group different copies of the same Nup together and then sort by name, adjacently numbered Nups are not necessarily spatially proximate. To optimize the visualization of the AE matrix, this study calculated the contact matrix between Nups based on the constricted state structural model and employed hierarchical clustering algorithms to rearrange the Nup order to better reflect spatial proximity relationships. In the AE matrix diagram, according to the set cutoff distance, the AE value regions can be clearly divided into three submatrices, corresponding to the three structural rings of the subunit: from left to right, the cytoplasmic ring, nuclear ring, and inner ring. The AE points in non-matrix regions represent Nups connecting different rings, namely Nup155-5 and Nup155-4.

Analysis results show that the overall AE values for the constricted-to-dilated state transition process are higher than those for the dilated-to-constricted process, reflecting that the former involves more significant structural changes. Additionally, AE values within the two outer rings are generally small and uniformly distributed, while AE values in the inner ring are overall larger with obvious fluctuations, containing multiple significantly high-value regions, indicating that the degree of structural change in the inner ring during NPC two-state transitions is markedly greater than in the outer rings. This conclusion is consistent with previous analysis results, so subsequent analysis will focus primarily on inner ring structure.

### Behaviors of inner-ring linker-Nup and FG-Nup in the two-state transition

The NPC scaffold structure contains multiple types of linker Nups, among which Nup155 directly connects the inner ring with the two outer rings, while other linker Nups mainly participate in connections within individual ring structures. Previous studies have confirmed that inner ring linker Nups regulate the relative positions of inner ring subunits by dynamically changing their connection conformations, thereby achieving inner ring contraction and expansion[7, 52, 53]. The simulation results of this study also validate this phenomenon. As shown in Figure 8, linker Nups primarily present linear conformations with their terminal regions binding to each other. They regulate the distance between inner ring subunits through hinge-like motion mechanisms, utilizing angular changes. In the AE matrix, inner ring linker Nups show relatively high AE value combinations in the inner ring region, mainly including: Nup35-3 with its neighboring nucleoporins, Nup35-2 with its neighboring nucleoporins, and Nup98-2 and Nup98-3. According to the definition of AE, changes in relative angles between these nucleoporins necessarily lead to increased AE values. The simulation results of this study show good consistency with existing research conclusions.

**Figure 6.**
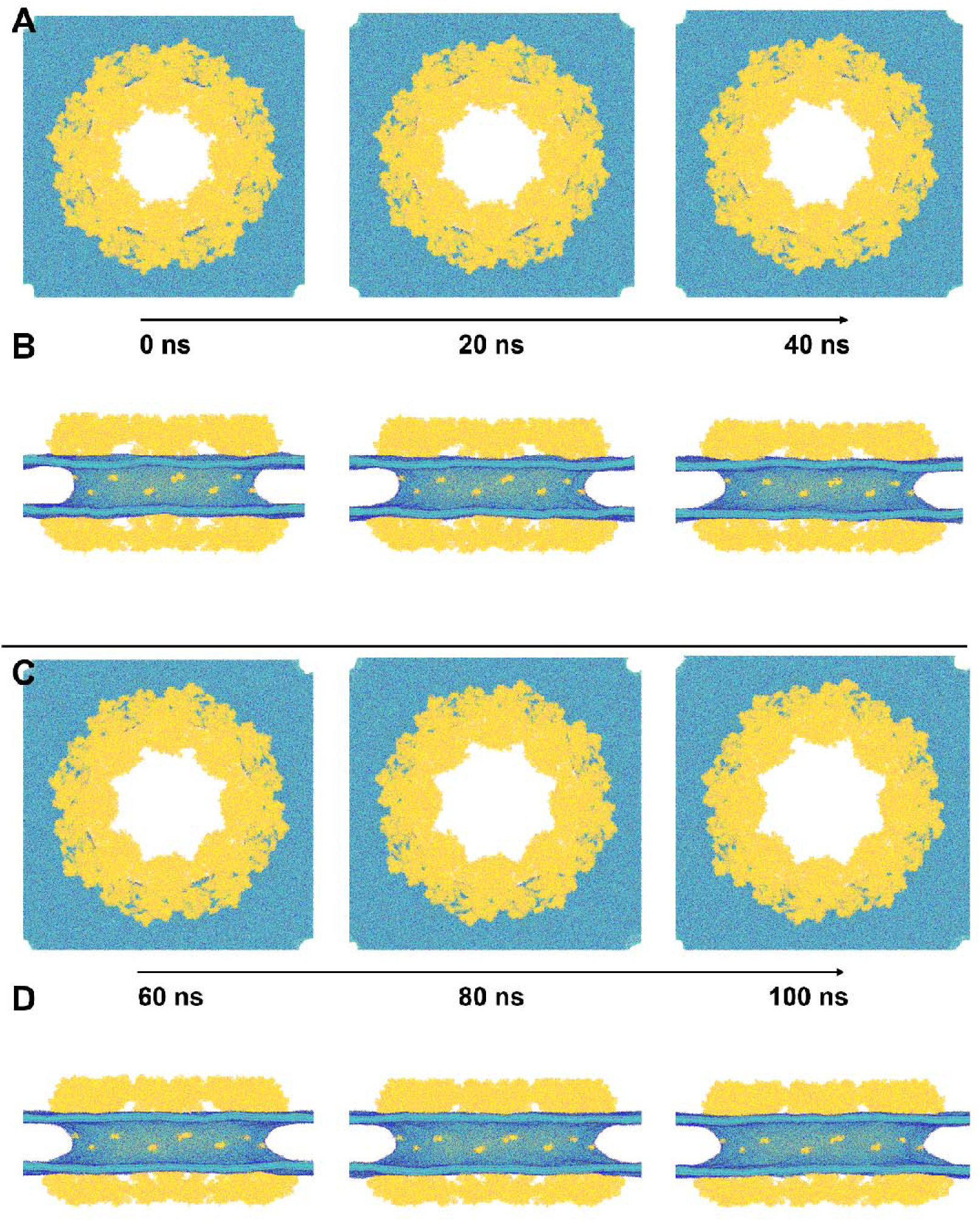
: Process of NPC scaffold structure transition from constricted to dilated state. Six snapshots were selected evenly from the 100 ns simulation process. Yellow represents the NPC scaffold structure, and blue represents the nuclear membrane. (A) System states at 0 ns, 20 ns, and 40 ns, respectively, top view. (B) System states at 0 ns, 20 ns, and 40 ns, respectively, side view. (C) System states at 60 ns, 80 ns, and 100 ns, respectively, top view. (D) System states at 60 ns, 80 ns, and 100 ns, respectively, side view.

**Figure 7.**
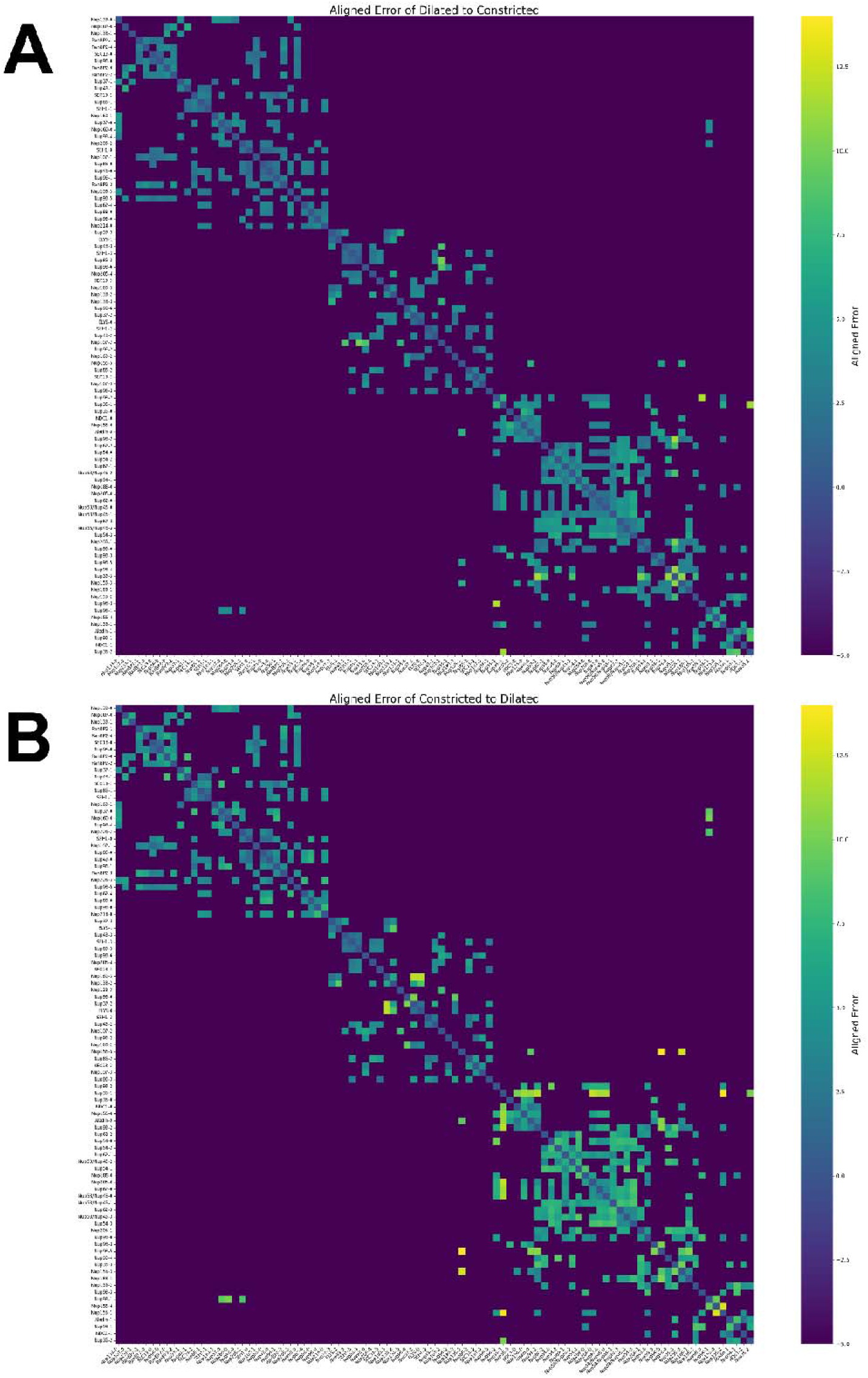
: AE matrix of initial and final conformations during NPC scaffold structure two-state transition. The left side and bottom are labeled with each Nup within the subunit of the NPC scaffold structure using "Nup name + copy number." For visualization purposes, AE values between two Nups filtered out by the cutoff distance in the matrix are uniformly marked as -5. AE values on the diagonal are 0, while all other AE values are greater than 0. (A) Dilated to constricted state. (B) Constricted to dilated state.

**Figure 8:**
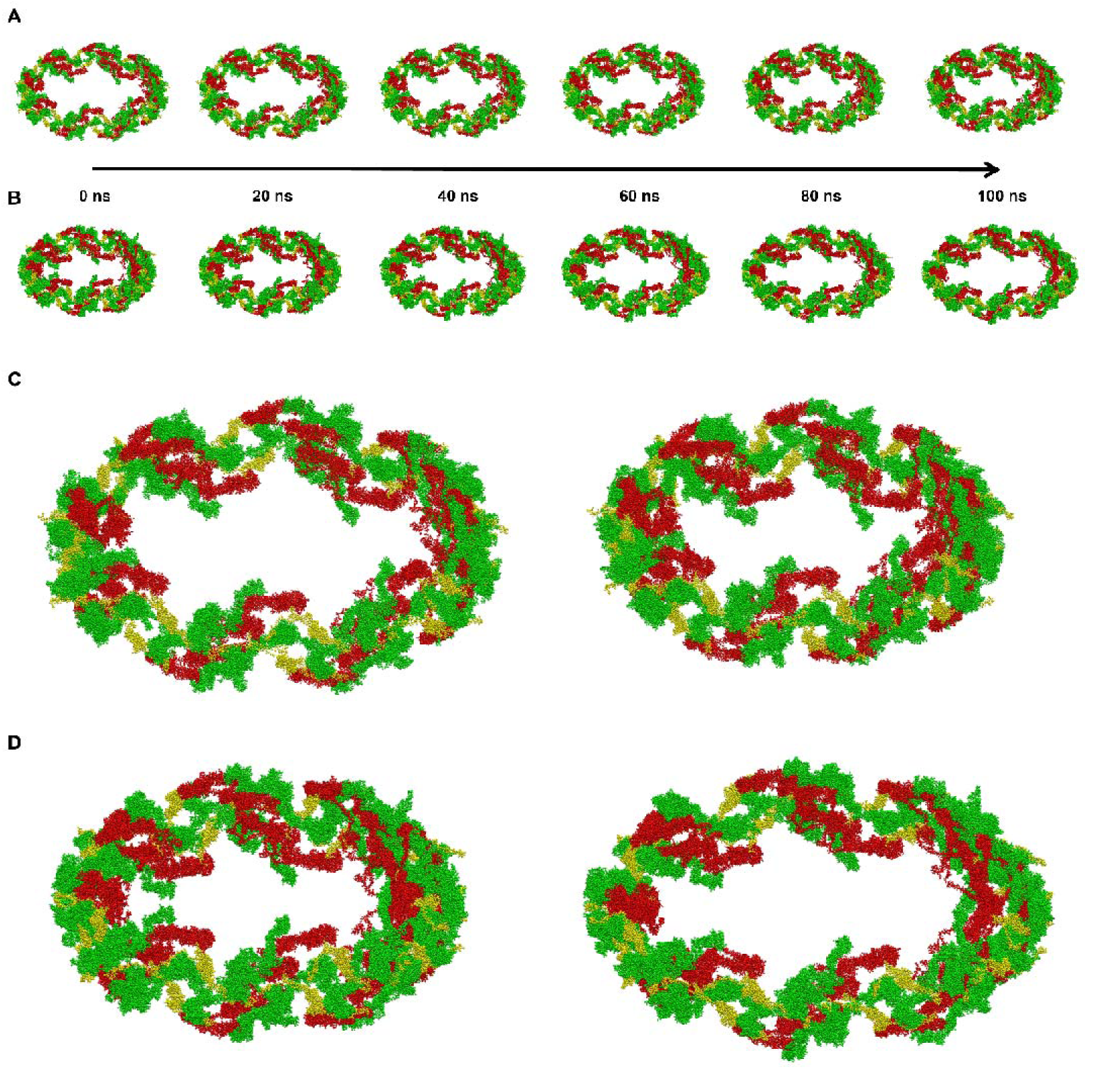
C**h**anges **in inner-ring linker-Nup during NPC scaffold structure two-state transition.** Yellow: Nup35-0, Nup35-1, Nup35-2, and Nup35-3. Red: Nup93-0, Nup93-1, Nup93-2, and Nup93-3. Green: Nup155-1, Nup155-2, Nup155-3, and Nup155-4. Six snapshots were selected evenly from the 100 ns simulation process. (A) Dilated to constricted state, oblique view. (B) Constricted to dilated state, oblique view. (C) Dilated to constricted state. Left: initial state; Right: final state. (D) Constricted to dilated state. Left: initial state; Right: final state.

During the two-state transition process, the AE matrix displays a significant feature: several Nups in the central region of the inner ring exhibit highly tight interactions, manifested as a continuous region of average AE values on the AE matrix. These Nups include Nup54-3, Nup58-3, Nup62-3, Nup58-1, Nup58-0, Nup62-0, Nup205-0, Nup188-0, Nup54-1, Nup58-2, Nup62-1, Nup54-2, Nup54-0, and Nup62-2 (in matrix numbering order). Notably, this region encompasses all inner ring FG-Nup copies while also including Nup205 and Nup188.

All FG-Nups in the inner ring are collectively referred to as the FG-Nup heterotrimer. As shown in Figure 9, these FG-Nups are tightly intertwined with each other, forming a regular "core" structure. This structure serves as the functional unit executing the core functions of the NPC scaffold, with its extending FG repeat sequences directly regulating nucleocytoplasmic material transport. During the two-state transition process, they maintain close contact throughout, with their overall conformation remaining stable and only undergoing minor changes in mutual spacing. This spacing change reflects the contraction or expansion of the inner ring. This dynamic process is jointly regulated by other inner ring Nups (including nuclear membrane-anchoring Nups and linker Nups): nuclear membrane-anchoring Nups transmit nuclear membrane morphological changes to the NPC scaffold structure, driving its expansion and contraction; simultaneously, inner ring linker Nups mediate relative displacement of inner ring subunits. This ultimately leads to changes in FG repeat sequence anchoring sites and nuclear pore channel diameter variations, potentially regulating the distribution of FG repeat sequences within the channel and achieving control over nucleocytoplasmic transport.

**Figure 9:**
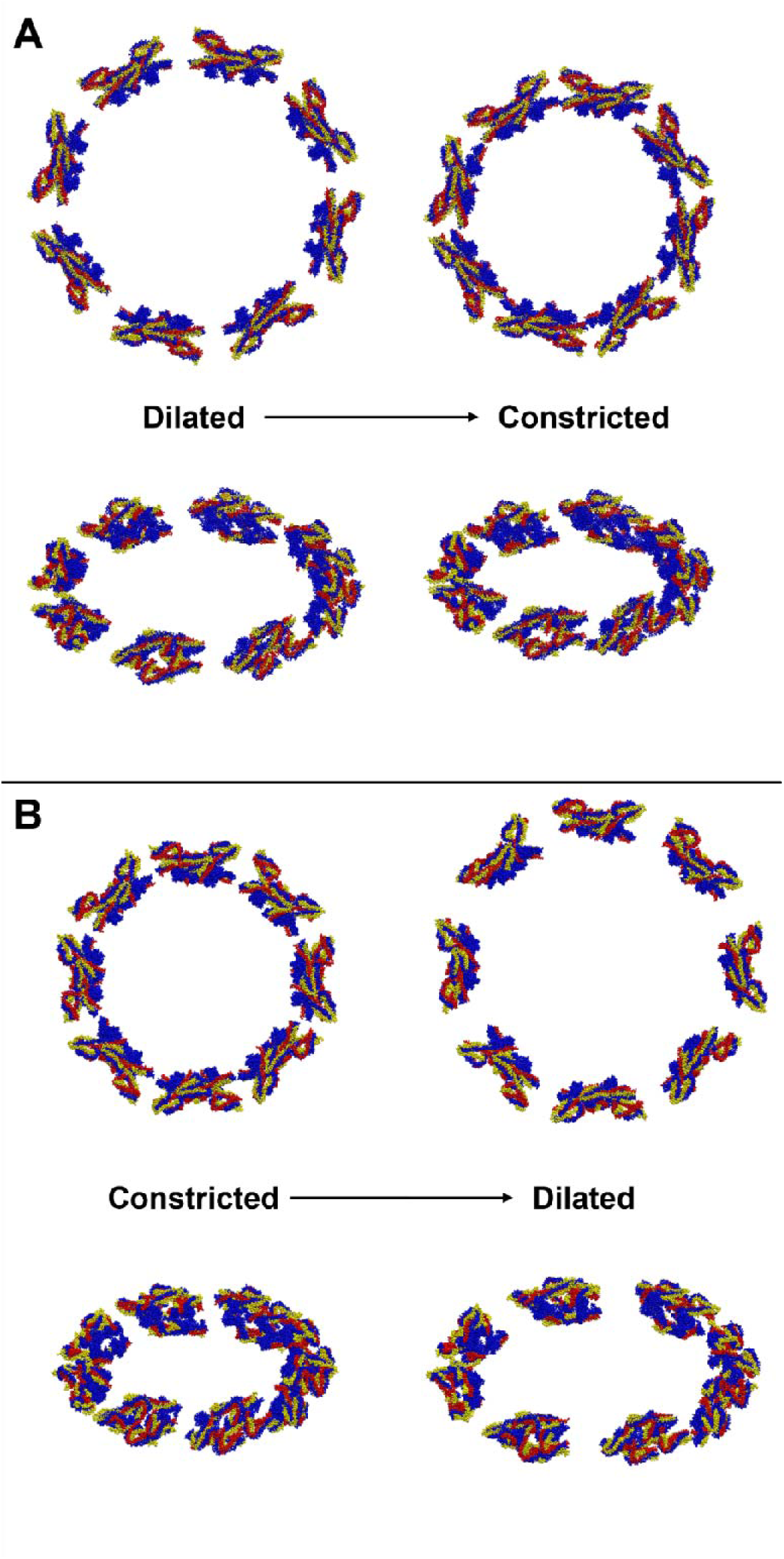
C**h**anges **in inner ring core during NPC scaffold structure two-state transition.** Blue: Nup54-0, Nup54-1, Nup54-2, and Nup54-3. Red: Nup58-0, Nup58-1, Nup58-2, and Nup58-3. Yellow: Nup62-0, Nup62-1, Nup62-2, and Nup62-3. (A) Dilated to constricted state. Left: initial state, Right: final state. Top: top view, Bottom: oblique view. (B) Constricted to dilated state. Left: initial state, Right: final state. Top: top view, Bottom: oblique view.

### Distribution of the inner-ring FG repeat sequences in nuclear pore

The left half of Figure 10 displays simulation snapshots after equilibration in the constricted state. In the left panel, as the restraint force constant decreases (from top to bottom), the distribution of FG repeat sequences within the nuclear pore gradually becomes more dispersed, with the "knot" in the central region gradually disappearing. When the force constant decreases to k = 0.001 kJ·mol_⁻_¹·nm_⁻_², the FG repeat sequences essentially exhibit an average distribution. In the right panel, the equilibrium conformational distribution of unrestrained FG repeat sequences is similar to the distribution at k = 0.001 kJ·mol_⁻_¹·nm_⁻_².

**Figure 10:**
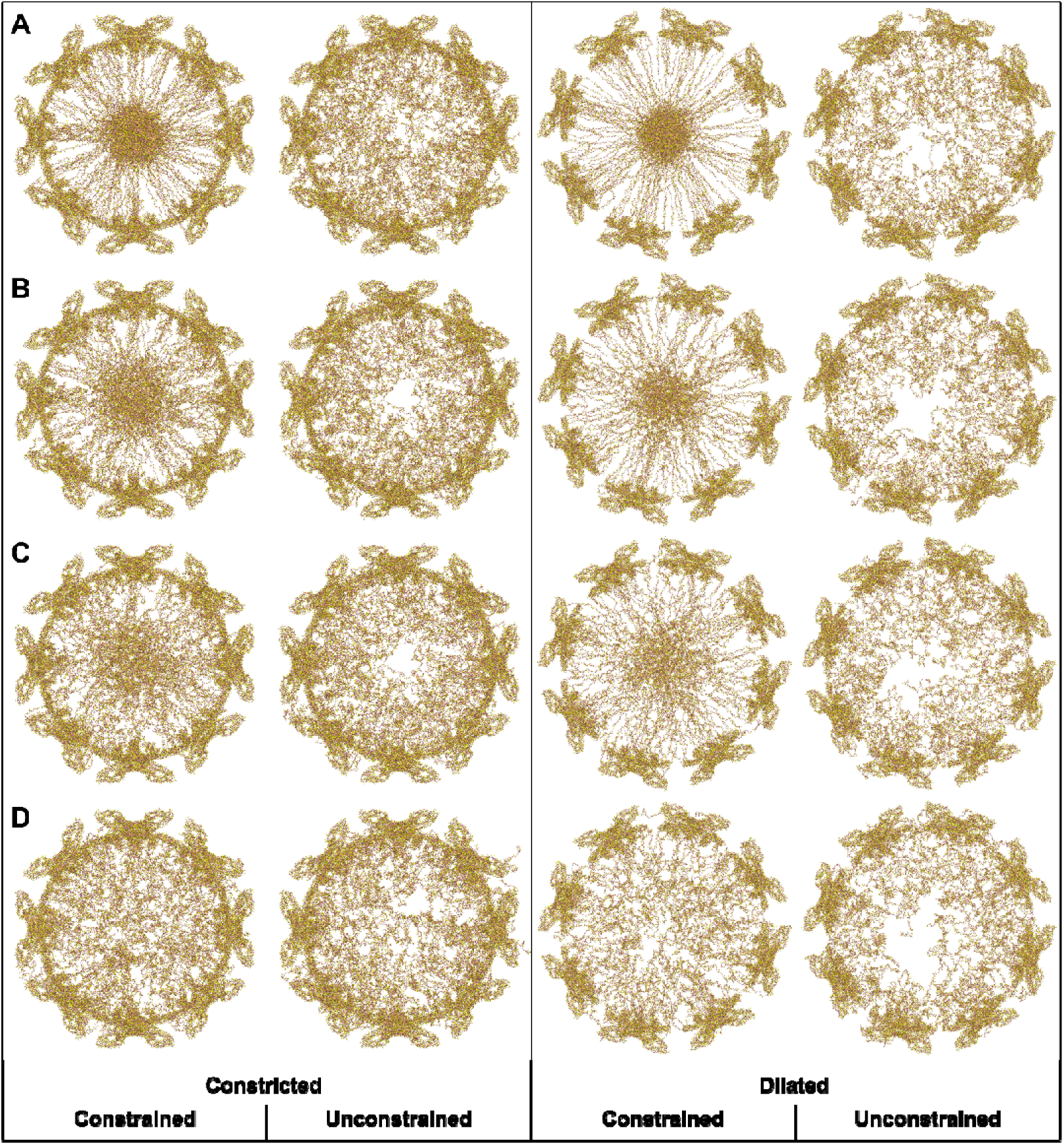
S**i**mulation **snapshots of NPC constricted and dilated state inner-ring FG-Nup system.** From left to right: constricted state restrained final conformation; constricted state unrestrained final conformation; dilated state restrained final conformation; dilated state unrestrained final conformation. From top to bottom, the restraint force constants are: (A) k = 0.05 kJ·mol_⁻_¹·nm_⁻_ ². (B) k = 0.01 kJ·mol_⁻_¹·nm_⁻_ ². (C) k = 0.005 kJ·mol_⁻_¹·nm_⁻_². (D) k = 0.001 kJ·mol_⁻_¹·nm_⁻_².

Similarly, the simulation snapshots after equilibration in the dilated state shown in the right half of Figure 10 demonstrate that the distribution of FG repeat sequences also gradually becomes more dispersed as the restraint force decreases, approaching uniform distribution at k = 0.001 kJ·mol_⁻_¹·nm_⁻_². Due to the larger nuclear pore channel diameter in the dilated state, its distribution density is relatively sparser compared to the constricted state. After removing the restraint force, the unrestrained distribution is also similar to that of the constricted state but with higher sparsity.

Figure 11 shows radial FG repeat sequence density distribution curves obtained from statistical analysis of the last 25% of trajectory frames. Panel A shows distribution curves for the restrained state, while Panel B shows distribution curves for the unrestrained state. Although individual distribution curves exhibit some randomness, they still present obvious regular characteristics.

**Figure 11:**
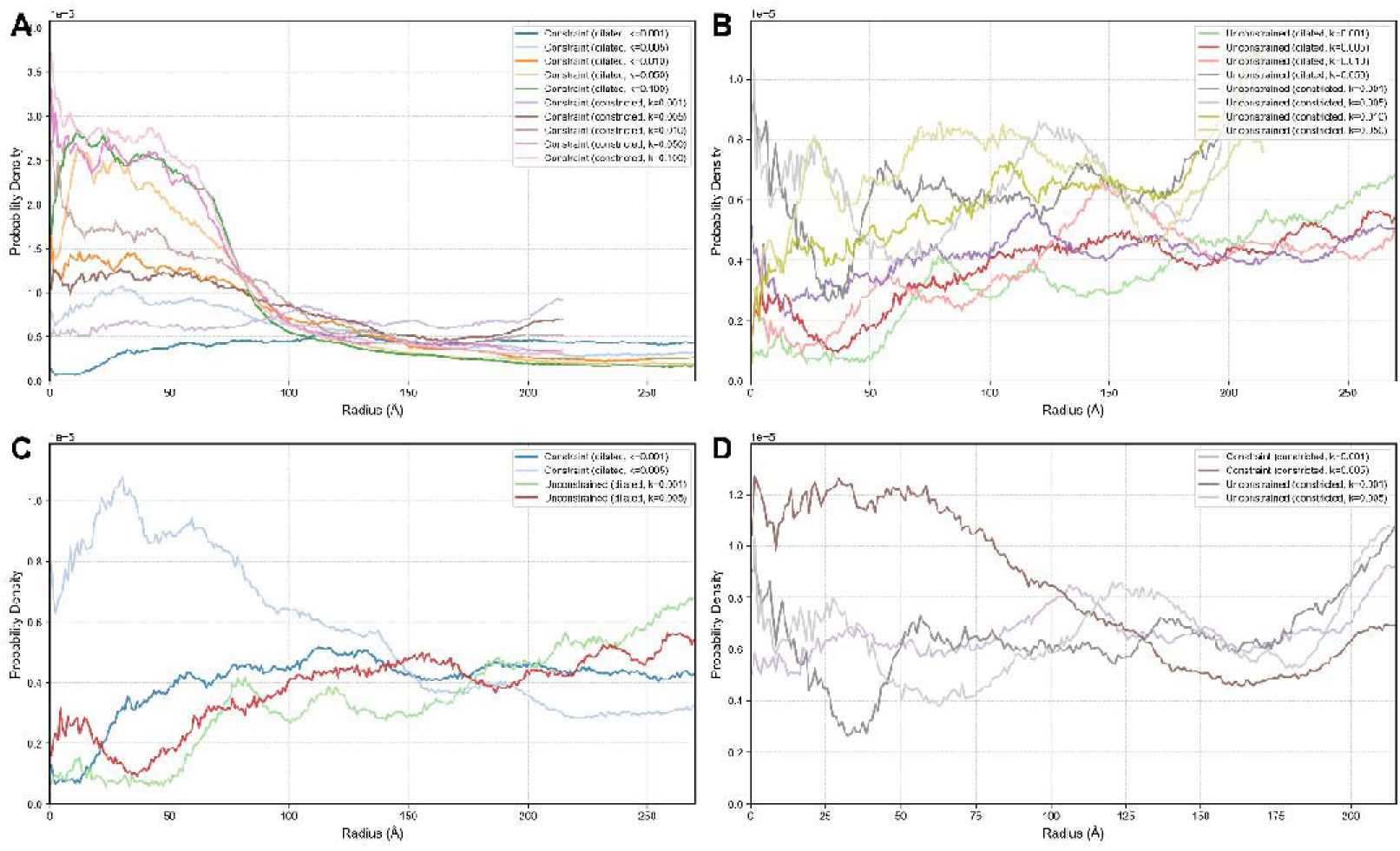
D**e**nsity **distribution of FG repeat sequences along nuclear pore radius.** The representative curve colors for each state remain consistent. (A) Restrained state NPC: comparison between constricted and dilated states. (B) Unrestrained state NPC: comparison between constricted and dilated states. (C) Dilated state NPC: comparison between restrained states with force constants k = 0.005 kJ·mol_⁻_¹·nm_⁻_² and k = 0.001 kJ·mol _⁻_ ¹·nm _⁻_ ² and the unrestrained state. (D) Constricted state NPC: comparison between restrained states with force constants k = 0.005 kJ·mol_⁻_¹·nm_⁻_² and k = 0.001 kJ·mol_⁻_¹·nm_⁻_² and the unrestrained state.

In restrained state simulations, the constricted and dilated states exhibit similar overall change trends. In the central vicinity region, the density distribution of FG repeat sequences shows obvious peaks, with these peaks increasing as the force constant increases. Under the same force constant conditions, due to the smaller nuclear pore channel cross-sectional area, the density distribution in the constricted state is consistently higher than in the dilated state. These observations are essentially consistent with the analysis conclusions from simulation snapshots. In unrestrained state simulations, the change trends for constricted and dilated states are essentially consistent, manifesting as slight increases with distance from the nuclear pore center, unaffected by the force constant magnitude in previous restrained state simulations. This corresponds to the central region void phenomenon observed in simulation snapshots. Additionally, the overall density distribution in the constricted state remains higher than in the dilated state.

As shown in Panels C and D of Figure 11, by comparing the FG repeat sequence density distributions between restrained and unrestrained states, when the restraint force constant is k = 0.001 kJ·mol_⁻_ ¹·nm_⁻_², its density distribution essentially matches that of the unrestrained state. Therefore, in subsequent simulations, k = 0.001 kJ·mol_⁻_ ¹·nm_⁻_ ² can be adopted as an approximate parameter for unrestrained state distribution, maintaining the distribution characteristics of FG repeat sequences while preventing their appearance in non-target regions.

## CONCLUSION

This study systematically elucidated the dynamic change mechanisms of human nuclear pore complex scaffold structure and its regulatory effects on FG repeat sequence distribution through a multi-scale computational approach combining elastic network models (ENM) and molecular dynamics (MD) simulations. ENM analysis successfully identified the first three low-frequency collective motion modes of the NPC scaffold structure, revealing the coupling relationship between inner ring radial contraction/expansion and outer ring axial displacement. MD simulations achieved dynamic conformational transitions between constricted and dilated NPC scaffold states, validating the motion modes predicted by ENM and obtaining detailed kinetic information of the two-state transition process, including nuclear membrane spacing changes, bending angle adjustments, and dynamic regulation of nuclear pore channel diameter. Aligned error analysis further confirmed that the degree of structural change in the inner ring is significantly greater than in the outer rings, highlighting the central role of the inner ring in NPC conformational transitions. Although ENM as a coarse-grained model has certain limitations, it can still effectively capture the main collective motion characteristics of NPC scaffold structure, providing meaningful structural biology insights for understanding nuclear pore complex conformational dynamics.

Based on the above multi-scale research results, this study proposes a possible nucleocytoplasmic transport regulatory pathway model: under physiological conditions, changes in nuclear membrane morphology are transmitted to the NPC scaffold structure through transmembrane nucleoporins, driving dynamic regulation of inter-subunit spacing in the inner ring[26]. Inner ring linker proteins (such as Nup35 and Nup98) mediate this process through hinge-like motion mechanisms, while the inner ring FG-Nup heterotrimer, serving as the core functional unit, undergoes corresponding spatial position changes with inner ring contraction/expansion while maintaining overall conformational stability. These positional changes directly affect the distribution of anchoring points for FG repeat sequences within the nuclear pore channel, thereby regulating FG barrier density. The outer rings (cytoplasmic ring and nuclear ring) influence the geometric configuration of the nuclear pore channel through axial coupling motion with the inner ring, and this configurational change may further affect the FG barrier. This model provides new support for the dilation model[19] theoretical framework, emphasizing the key role of NPC scaffold structure dynamics in nucleocytoplasmic transport regulation.

This study still has some limitations, mainly reflected in the following aspects: First, although the periodic boundary condition optimization algorithm used in ENM analysis significantly reduced computational complexity, it may introduce certain systematic errors, particularly in the calculation of XY plane eigenvector components; Second, although the biasing potential method used in MD simulations can achieve two-state transitions, it may not fully reflect the real transition pathways under physiological conditions[26]; Additionally, the simulation of FG repeat sequences employed simplified coarse-grained models, which may not adequately capture their complex dynamic behavior and interactions with transport receptors. Future research should consider adopting more refined all-atom simulation methods to perform full-system simulations including complete NPC structure and FG repeat sequences, directly validating the possible nuclear pore functional regulatory mechanisms proposed in this study, and providing theoretical guidance for the treatment of related diseases and drug design.

## ACKNOWLEDGMENTS

This work is supported by the National Key Research and Development Program of China (2021YFA1301504), the National Natural Science Foundation of China (91953101), and the Chinese Academy of Sciences Strategic Priority Research Program (XDB37040202).

## CONFLICT OF INTEREST

There is no conflict of interest to report.

## Notes

### Competing Interest Statement

The authors have declared no competing interest.

